# Estimating the evolution of human life history traits in age-structured populations

**DOI:** 10.1101/002584

**Authors:** Ryan Baldini

## Abstract

I propose a method that estimates the selection response of all vital rates in an age-structured population. I assume that vital rates are determined by the additive genetic contributions of many loci. The method uses all relatedness information in the sample to inform its estimates of genetic parameters, via an MCMC Bayesian framework. One can use the results to estimate the selection response of any life history trait that is a function of the vital rates, including the age at first reproduction, total lifetime fertility, survival to adulthood, and others. This method closely ties the empirical analysis of life history evolution to dynamically complete models of natural selection, and therefore enjoys some theoretical advantages over other methods. I demonstrate the method on a simulated model of evolution with two age classes. Finally I discuss how the method can be extended to more complicated cases.

## 1 Introduction

Because evolution is a multigenerational process, multigenerational data provide ideal information with which to test evolutionary hypotheses. Studies using multigenerational human data are increasingly common (Stearns et al., 2010), but the methodology varies widely, with little commentary on which is best. The complexity of evolution in recent human history - which, among other things, involves overlapping generations, rapid demographic and environmental change, and cultural transmission of behaviors - makes its analysis very difficult.

Here I propose a method in which dynamically complete evolutionary models are fit directly to empirical data, so that theory and methods are as closely aligned as possible. This allows us to infer the results of selection across generations (at least in the short term). In this paper I focus particularly on the evolution of life history characteristics in age-structured populations, as all human populations involve age structure. The idea is to estimate the genetic variation underlying all vital rates, given some model of genetic inheritance. From these estimates and the model underlying them, the selection response of almost any life history characters can be estimated. I use a Bayesian MCMC approach to estimate the genetic parameters using all relatedness information in the population. I demonstrate the general method on simulated data with two age classes and quantitative-genetic inheritance. Later I discuss how the method can be applied to more complex cases.

## 2 The method

The method I propose is perhaps the most straightforward application of evolutionary theory to multigenerational data. We simply propose a dynamically complete model of evolution (likely from the population genetics literature), derive its state variables^1^, and estimate those variables from the real data. This allows us to understand how forces have changed each variable - and, if the model is accurate and conditions don’t change, allows prediction for future generations.

I focus here on evolution in an age-structured population. Genetic models of age-structured selection (Charlesworth, 1994) show that genetic variation in vital rates - the rates of survival and fertility at each age for each genotype - determine the dynamics of selection. The most direct application of such models is, therefore, to estimate the genetic variation of each vital rate in the population. Most life history traits are functions of these vital rates - including the expected age at first reproduction, lifetime offspring count, lifespan, and so on. We can therefore derive estimates of the evolution of any such life history trait from those of the vital rates.

To have a dynamically complete model requires that we specify an explicit model of inheritance. In principle, we can fit any genetic model we like, so long as there is sufficient data to identify all parameters. In the example I provide below, I assume a Normal distribution of additive genetic values (or “breeding values”) for all traits, with a constant covariance structure. This assumption is impossible to justify exactly under selection (Turelli, 1988; Turelli and Barton, 1990), but it is a common and accurate approximation for short-term evolution when many loci contribute to the traits under evolution (Turelli and Barton, 1994).

I use a Bayesian MCMC framework to estimate each individual’s breeding value. These estimates, though imprecise for each individual, can collectively produce accurate estimates of genetic means and covariances for the entire population. Estimates of individuals’ breeding values are informed by their own reproductive outcomes and those of their relatives, in a way that is consistent with the model (see the example below). Thus, the method requires multigenerational datasets with complete reproductive histories of each individual, identifying each individual’s parents in the data (if they exist).

## 3 An example with two age classes

### 3.1 The model

Consider a population of sexually-reproducing organisms that are classified into two age classes: ages 1 and 2. Each genotype is specified by its breeding values for three vital rates. These are *M*_1_ and *M*_2_, a genotype’s expected number of offspring that survive to the next census time for ages 1 and 2, respectively (this includes both fertility and newborn survival); and *P*_2_, the probability that an individual of the genotype survives from age 1 to 2. Because reproduction occurs in pairs, achieved fertility may depend not only on one’s own age and genotype, but also those of the mate. Thus, *M*_1_ and *M*_2_ cannot be a fixed property of the genotype, as they depend in general on the state of the population. Other models (e.g. Charlesworth 1994) assume that fertility is solely determined by the female, from which fixed values of *M* can be derived. This is unrealistic for humans, so I assume an averaging process instead: for any pair of ages *x* and *y*, the pair’s expected fertility, *M* (*x*, *y*), is the mean of each genotypes’ *partial* contribution to fertility, denoted by *M̂*:

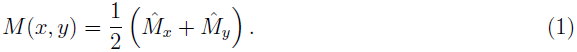

Each *M̂* corresponds to the genotype of one individual in the pair. Thus, this model assumes that men and women determine fertility equally.

The relevant vital rates for each genotype, then, are *M̂*_1_, *M̂*_2_, and *P*_2_. The breeding values for these vital rates cannot be normally distributed, however, as all are constrained to be positive, and the probability of survival must lie in the interval [0,1]. I therefore represent genotype by a transformation of these vital rates which produces a vector **g**, each element of which is defined on the real number line. Thus, each genotype has a characteristic vector **g** = (*g*_1_, *g*_2_, *g*_3_), which correspond to *M̂*_1_, *M̂*_2_, and *P*_2_, respectively, as defined below.

Because human fertility for any short age class cannot be very large, I impose an upperlimit on *M̂*. I assume that the achieved fertility of a pair is binomially distributed with a maximum of *k*. Specifically, if *f* is the achieved fertility of a pair of ages *x* and *y* ∈ {1,2}, then

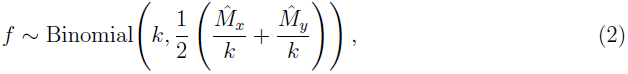

where, for ages 1 and 2,

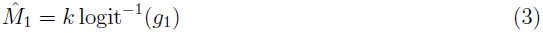

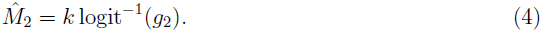

This just says that the fertility of a pair whose parents are of age *x* and *y* is binomially distribution with probability parameter equal to the mean of the inverse-logit^2^ of the breeding values *g*_*x*_ and *g*_*y*_. The parameterization ensures that the pair’s expected fertility satisfies equation (1). In this example, I choose *k* = 3. Neither this value nor the binomial distribution is required for the method; other discrete distributions, e.g. the Poisson, could be used as well.

For survival, I assume

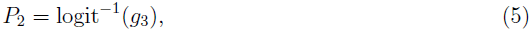

i.e. *g*_3_ corresponds to *P*_2_ via an inverse-logit transformation. For observed data, let *s* be the outcome for whether an individual survives or not. Then *s* is a Bernoulli random variable:

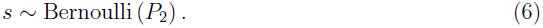

Because the inverse-logit function’s domain is the entire real number line, we are free to assume that

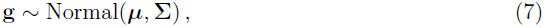

where ***µ*** is the vector of means for each *g*, and **Σ** is their covariance matrix. These are, ultimately, the state variables of the model. Equation (7) is the probability model for individuals without parents in the data. For individuals whose parents data are provided, the assumption of random mating and constant covariances implies that **g**_*o*_, the breeding values of a pair’s offspring, are distributed as

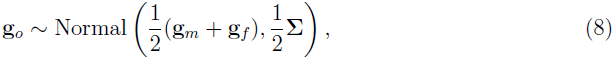

where **g**_*m*_ and **g**_*f*_ are the breeding values of the mother and father, respectively.

Table 1 reviews all terms in the model.

**Table 1.** Description of terms in the model

| Variable | Description |
| --- | --- |
| $\hat{M}_1$ | A genotype's partial contribution to a pair's fertility at age 1, via equation(1) |
| $\hat{M}_2$ | A genotype's partial contribution to a pair's fertility at age 2, via equation(1) |
| $P_2$ | A genotype's probability of surviving to age 2 |
| $g_1$ | A genotype's breeding value corresponding to $\hat{M}_1$, via equation (3) |
| $g_2$ | A genotype's breeding value corresponding to $\hat{M}_2$, via equation (4) |
| $g_3$ | A genotype's breeding value corresponding to $P_2$, via equation (5) |
| $\mathbf{g}$ | The vector $(g_1, g_2, g_3)$ specifying a complete genotype |
| $\boldsymbol{\mu}$ | The vector $(\mu_1, \mu_2, \mu_3)$ specifying the means for $\mathbf{g}$ |
| $\boldsymbol{\Sigma}$ | The covariance matrix for $\mathbf{g}$ |

### 3.2 Fitting the model

The method proceeds by fitting (2), (6), (7), and (8) to the data. This is sufficient because the above equations completely define the model. For individuals whose parents are not present, we use (7); otherwise, parental information is used as in (8). Model (2) is fit to the fertility of each pair at each age, and (6) is fit to the survival outcome of each individual at every time period.

If each person’s parents are identified in the data, then the method uses the information from every relative to estimate each individual’s genotype. There is no need to include relatives other than the parents in (8). The logic for this is as follows. Suppose that an individual’s sibling is in the dataset. The sibling’s outcomes correlate with ego’s outcomes only insofar as they receive identical genes from the parents. Thus, high sibling fertility at age 1 therefore increases the likelihood that the sibling has a large *g*_1_, suggesting that the parent has a large *g*_1_, ultimately suggesting that ego has a large *g*_1_. Similar logic applies for cousins, grand-offspring, and so forth. The strength of the information decays geometrically with decreased relatedness, but the model accounts for all of it.

I fit the model by sampling 5000 individuals of age 1 to produce an initial cohort, from which all later individuals are birthed. I use five total generations below. The initial cohort is sampled from (7), thereby assuming that they are unrelated. In real datasets, the initial cohort may include close relatives; the model is easily adjusted to account for this information.

### 3.3 Estimates for genetic parameters

Table 2 shows the estimates for the distribution of **g** from the simulated data. The 95% interval estimates overlap the true values in each case. The estimates correctly suggest that the genetic covariances between all vital rates are negative, although the 95% interval estimate overlaps 0 for *C*_2_3. Of course, the accuracy of the estimates depends on the quantity of the data. Inferring both genetic and selective effects requires are large sample size. In most cases, the method cannot be expected to give accurate estimates when sample sizes are less than ~1000.

**Table 2.** Point estimate is the median of the posterior. The interval estimate is the 95% highest-probability-density interval. *V*_*i*_ refers to the variance of *g*_*i*_; *C*_*ij*_ refers to the covariance between *g*_*i*_ and *g*_*j*_; all are elements of the matrix **Σ**. *N* = 14105, which descend from an original cohort of size 5000.

| variable | point estimate | interval estimate | true value |
| --- | --- | --- | --- |
| $\mu_1$ | -0.99 | (-1.05, -0.92) | -1 |
| $\mu_2$ | -0.04 | (-0.11, 0.03) | 0 |
| $\mu_3$ | 0.98 | (0.94, 1.03) | 1 |
| $V_1$ | 0.97 | (0.78, 1.21) | 1 |
| $V_2$ | 0.96 | (0.67, 1.26) | 1 |
| $V_3$ | 0.09 | (0.02, 0.22) | 0.1 |
| $C_{12}$ | -0.28 | (-0.43, -0.12) | -0.3 |
| $C_{13}$ | -0.09 | (-0.17, -0.01) | -0.1 |
| $C_{23}$ | -0.06 | (-0.21, 0.05) | -0.1 |

### 3.4 Estimates for selection response of vital rates

Charlesworth (1994) shows that for an additive, quantitative genetic trait *g*, the response to weak selection over one time period can be approximated by

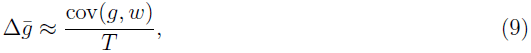

where *w* is relative fitness for genotype *g*, and *T* is a measure of generation time. The exact form of *w* and *T* depend on the details of the model. I derive these quantities in the appendix for the model used here. As is common, I use *m̂*_*x*_, the partial number of *daughters* produced by a genotype at age *x*, instead of *M̂*_*x*_. Under even sex ratios, *m̂* = *M̂*/2. The fitness formula is then

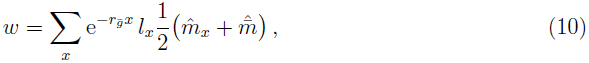

where *l*_*x*_ is the probability of surviving to age *x* (recall *l*_1_ = 1, so *l*_2_ = *P*_2_) and 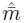 is the mean partial number of daughters in the population, given by

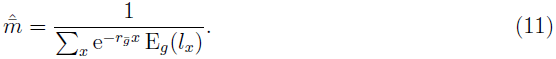

*E*_*g*_() denotes expectation over the entire distribution of genotypes. The term *r*_*ḡ*_ is a measure of the growth rate of the population implicitly defined by

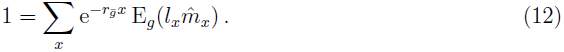

Finally,

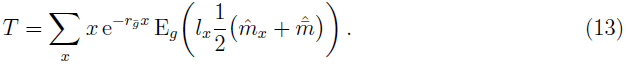

Equation (10) is a frequency-dependent measure of fitness (i.e. its value for any genotype depend on the state of the population) that is appropriate for the present model. It differs somewhat from measures provided elsewhere (Charlesworth, 1994) because most derivations either assume asexual reproduction or female-determined fertility. My model assumes shared fertility among pairs, which, together with random mating across age and genotype, implies that a genotype’s realized fertility is pulled toward the mean as in equation (10).

Table 3 shows the results. The available data correctly suggest, with 95% confidence, that *μ*_1_ and *μ*_2_ will increase. The predicted directional change of *μ*_3_ is equivocal - presumably because the actual change is so close to zero. In each case, though, the true value lies within the 95% interval estimate. Because these traits all are positive components of fitness, the direct selection differential on each is positive. Thus, *μ*_3_ only decreases because *g*_3_ is negatively correlated with the other fertility traits, which show greater genetic variation.

**Table 3.** Point estimate is the median of the posterior. The interval estimate is the 95% highest-probability-density interval. *N* = 14105, which descend from an original cohort of size 5000.

| variable | point estimate | interval estimate | actual value |
| --- | --- | --- | --- |
| $r_{\bar{g}}$ | -0.008 | (-0.021, 0.005) | -0.003 |
| $T$ | 1.480 | (1.473, 1.487) | 1.482 |
| $\Delta\mu_1$ | 0.059 | (0.043, 0.081) | 0.058 |
| $\Delta\mu_2$ | 0.043 | (0.021, 0.067) | 0.040 |
| $\Delta\mu_3$ | -0.005 | (-0.014, 0.006) | -0.008 |

### 3.5 Estimates for selection response of composite life history characters

Most life history traits are functions of the vital rates treated in the model. A genotype’s expected lifespan, for example, is simply 1 + *P*_2_ in this model (recall that we have only accounted for individuals who have already survived to the first age class). The expected lifetime number of offspring to an individual who survives to age 2 is 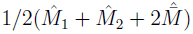. Thus, it is possible to determine the selection response of such traits from the changes in the vital rates themselves. Most such traits, however, are a non-linear function of the **g**, so that equation (9) is no longer valid. The simplest recourse is to take the difference between the means across one time step, as predicted by the model itself. These are readily produced by the above estimates.

Table 4 shows estimated selection response for expected lifespan, age at first reproduction, and lifetime number of offspring (see the caption for variable explanations). The method correctly predicts the direction of evolution for age of first reproduction and total number of offspring, with 95% confidence. Again, the interval estimates contain the true value in each case. We infer that selection is leading to slightly earlier reproduction and a greater number of total offspring per parent, but that lifespan may be slightly decreasing.

**Table 4.**
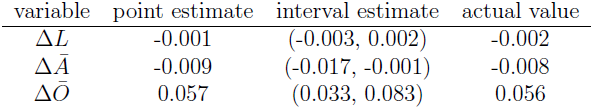
*L* is the mean lifespan among newborns. *A* is the mean age at first reproduction among those who survive to age 2. *O* is the expected number of offspring among those who survive to age 2. The point estimate is the median of the posterior. The interval estimate is the 95% highest-probability-density interval. *N* = 14, 105, which descend from an original cohort of size 5000.

| variable | point estimate | interval estimate | actual value |
| --- | --- | --- | --- |
| $\Delta \bar{L}$ | -0.001 | (-0.003, 0.002) | -0.002 |
| $\Delta \bar{A}$ | -0.009 | (-0.017, -0.001) | -0.008 |
| $\Delta \bar{O}$ | 0.057 | (0.033, 0.083) | 0.056 |

## 4 Fitting more realistic models

Two discrete age classes is clearly unrealistic for humans. In demographic studies, humans are usually broken into age classes of 5 or 10 years in length. As age classes become more numerous, the inference problem becomes more cumbersome, though not any more conceptually difficult. With 10 age classes, for example, we would need to estimate 20 parameters per person to fit the model in this paper. One way to reduce the dimensionality of the model, then, would be to assume a reproductive function that specifies a fertility course over the lifetime with just a few parameters. This method would have an inferential advantage in that the reproductive outcomes of each age provide information on the breeding values of every other age, because each realization informs the entire function. Another possible simplification of the method is to use continuous time, rather than discrete age classes. Continuous fertility and mortality functions are necessary in this case.

One major assumption in this paper’s model is that of a constant environment. That is, the expected survival and fertility of some genotype **g** is the same for all generations. This assumption cannot be true for most historical human populations; social, economic, and demographic conditions have changed enormously. Thus, it will be necessary to fit a model of temporal variation to the data. For the model of this paper, one would include a temporal effect in equations (3), (4), and (5). Reproductive outcomes can change over time because of environmental change or because of evolution in breeding values. The method allows us to separately estimate the effects of both because they are separated in the model.

Many have suggested that much human reproductive behavior is culturally transmitted - that is, learned socially from others (Cavalli-Sforza and Feldman, 1981; Boyd and Richerson, 1985). If this is the case, then any model based on vertically transmitted genes will be inappropriate. One can, in principle, start over with a cultural model, but this may require very different kinds of information (e.g. social networks) that are not readily available from historical data. Fortunately, if culture happens to be transmitted largely from parent to offspring, then the dynamics will be similar to the quantitative genetic example of this paper; we would still be able to infer cultural contributions to evolution from correlations between relatives.

Ultimately, if the model of choice is inaccurate, then inferences based on the model will not be accurate either. Probably the best method for any dataset is to fit many models and use a model comparison metric (e.g. AIC) to choose between them, or average their results. For each model, we derive the dynamical consequences, identify the parameters that need to be estimated, and use the data to estimate them as completely as the data allows.

## 5 Advantages of the method

### 5.1 Closely aligning empirics with theory

Evolutionary theory since Darwin has progressed mainly through the construction of mathematical models. The most satisfying models of evolution are dynamically complete: they specify how selection and other forces change the distribution of heritable variation through time. Most theory historically assumes that genes are the source of heredity, although alternative forms of inheritance (e.g. culture) have been analyzed as well. An ideal empirical study, then, would seek to fit these complete models to real data, thereby aligning the empirical and the theoretical as closely as possible.

This is precisely what the present method attempts to do. In general, the method is this: propose a model (or a collection of models) that might capture the evolution of some trait in a population, identify the state variables of the model, and estimate these variables for the population. In this paper’s example, I assumed an age-structured model of natural selection with a particular form of mating, fertility determination, and quantitative genetic structure. From these assumptions, I derived (following Charlesworth, 1994) that the state variables of the system, with regard to life history evolution, are the means and covariances of breeding values for age-specific survival and fertility rates. These are the variables that theory tells us to estimate. Fortunately, most life history traits are functions of these basic vital rates, so it is ultimately possible to estimate the dynamics of higher-level traits as well.

I am not aware of any study that attempts to fit a dynamically complete model of human life history evolution to real data. Most focus instead estimating the relationship between a few life history traits on some measure of fitness, which may not correspond to any particular model of selection. These studies either fail to specify an explicit model at all, or fail to estimate the state variables of the model. Thus, inferring evolutionary dynamics from such studies is impossible. These studies are still quite useful, but more complete analyses are within our grasp.

### 5.2 Consistent fitness measures

Past attempts to measure selection in human populations have struggled with the use of the fitness concept. Models of evolution in age-structured populations, for example, show that selection acts not just on the number of offspring produced, but also on the timing of reproduction over the lifetime. Thus, fitness measures like lifetime reproductive success (LRS) are technically incorrect. Researchers sometimes continue to use LRS in hope that it is a good approximation of more sophisticated measures of fitness (Stearns et al., 2010). It is not yet clear when this is a safe assumption for humans. Käär and Jokela (1998) showed for three premodern populations that individual measures of fitness based on LRS and one that is sensitive to reproductive timing (from McGraw and Caswell, 1996; see below) do not rank individuals equivalently, but that the two measures mostly agree for older females (double check this). Goodman et al. (2012) showed that LRS strongly predicted one’s contribution to the population four generations later in a Swedish population. Jones and Bliege Bird (2014) similarly showed that LRS has a positive but nonlinear relationship to the same fitness measure as in Käär and Jokela (1998) for a Utah population. In any case, a more sophisticated fitness measure should be preferred whenever possible.

Among studies that do use age-sensitive fitness, however, the assumed measure of fitness varies. Usually fitness is chosen without specifying any particular model of evolution. For example, McGraw and Caswell (1996) suggest a fitness measure from “philosophical underpinnings,” without showing how this quantity refers to the evolutionary process of gene frequency changes through time. The method uses individual-level realizations of vital rates to construct a Leslie matrix from which one can compute the dominant eigenvalue; the implication is that an individual’s fitness is the rate of growth of a population consisting solely of people that have the same survival and reproduction schedule as the individual in question. Käär and Jokela (1998), Korpelainen (2003) Pettay et al. (2005), Jones and Bliege Bird (2014), and probably others, have applied this method to human populations. Voland (1990) and Goodman et al. (2012) use a multi-generational approach: one’s fitness is taken to be the total number of one’s extant descendants, weighted by their relatedness to the individual and by the reproductive value of their age and sex. It is not clear what evolutionary process this corresponds to. Moorad (2012) calculates a fitness measure that, like McGraw and Caswells’, uses individual reproductive histories, but imputes vital rates for individuals who die before finishing reproduction in a possibly arbitrary way, and similarly fails to show how these relate to any fully dynamic evolutionary model. Without the connection to evolutionary models, we have no way of knowing which method is best.

I take the view that fitness is a model-specific concept; it can only be correct within the context of a model. The present method helps to resolve the fitness problem, then, because our model choice uniquely determines the correct fitness measure (if it exists). That is, one can derive the correct fitness from the mathematics of the explicit model which must be assumed for the method’s use. For the example analyzed in this paper, I derived an approximate measure of fitness under weak selection in the same way as Charlesworth (1994) but for a different fertility scheme. As a result, the exact fitness quantity of my model differs from Charlesworth’s.

Working directly with genetic variation also removes the conceptual difficulty of defining fitness at the individual level (see discussion McGraw and Caswell, 1996). In the genetic models of evolution in age-structured populations, the vital rates of *genotypes*, not individuals, determine the dynamics of selection. It is the frequencies of genes, after all, that evolve; an individual outcome is just one realization of the vital rate of its genotype. Thus, a measure of fitness at the individual level is not necessary. The empirical literature, however, seems to be occupied with the use of individual-level fitness. Each of the fitness measures proposed by the researchers mentioned above, for example, apply at the level of the individual. Since my method works directly with genetic strategies, defining and measuring individual-level fitness is unnecessary.

I do not claim that an individual-level fitness concept is impossible or incoherent. The vital rates of genotypes, after all, are estimated by averaging the reproductive outcomes of individuals with the genotype. Thus, a natural definition of individual fitness is one’s particular realization of the genotypic fitness quantity. For the model in this paper, individual fitness would be analogous to equation (10), where *m*(*x*) is half one’s achieved number of offspring at age *x*, and *l*(*x*) is a binary variable representing whether or not the individual survived to age *x*. The justification of this definition is that its average across all individuals of a single genotype equals that genotype’s fitness; it therefore corresponds directly to a dynamically complete evolutionary model. This definition differs from both McGraw and Caswell’s (1996) eigenvalue approach and from Moorad’s (2012) imputation approach, although it is similar to both.

## A Response to selection and a measure of fitness

The biparental determination of fertility in my model is different than in other treatments of age-structured selection. Charlesworth (1994), for example, assumes fertility is solely determined by the mother, and that the mating success of males is proportional to that of females for any genotype. I follow the methods of Charlesworth (1994) to derive here the appropriate (though still approximate) measure of fitness for the mating model in this paper. I start by deriving the results for a single locus with an arbitrary number of alleles; additivity across independent loci then allows extrapolation to multiple loci (see Charlesworth, 1994, pp. 173-177).

I assume that males and females are equal in numbers as zygotes, and share the same survival and reproductive schedule. This allows us to simplify the problem, because then the gene frequencies in males and females are the same for all ages (see Charlesworth, 1994, pp. 109-110 for details). It is convenient, then, to consider evolution only among maternally derived gametes; males will have the same results. Thus, let *p*_*i*_(*t*) be the frequency of allele *i* among zygotes derived from mothers at time *t*. Let *B*(*t*) be the total number of zygotes at time *t*. Let *N*_*ij*_(*x*, *t*) be the number of females with maternally inherited allele *i* and paternally inherited allele *j* of age *x* at time *t*. Let *M*_*ij*_(*x*, *t*) be the expected number of offspring produced by a female of genotype *ij* at age *x*. Then

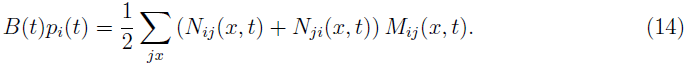

Random mating implies that 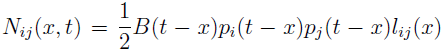, where *l*_*ij*_(*x*) = ∏_*y<x*_ *P*_*ij*_(*y*) is the probability that genotype *ij* survives to age *x*. Thus

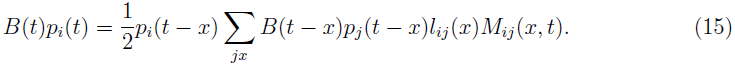

From the assumption of shared fertility determination within a pair, and random mating across ages and genotypes,

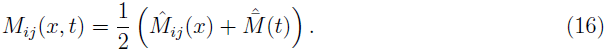

where 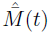 depends on the state of the population at time *t*. It is

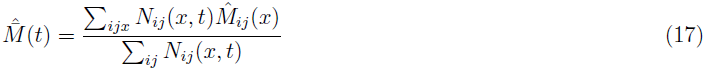

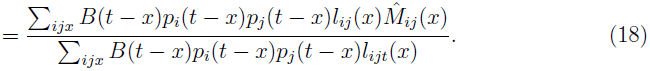

These equations specify the dynamical system. From them we can derive an approximation of Δ*p*_*i*_ under weak selection. The method follows Charlesworth (1994); see pp. 137-140 for details. The logic of the approximation is that, under sufficiently weak selection, each genotype approximately approaches a stable age distribution on a faster timescale than a change in *p*. Thus *B*(*t* – *x*)/*B*(*t*) can be approximated by *e*^−*r*_*p̄*_*x*^, with *r*_*p̄*_ being a measure of population growth rate as defined below. We ultimately find

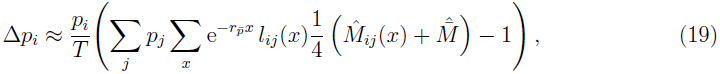

where

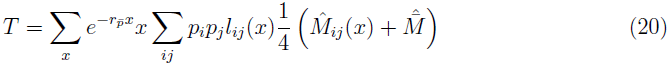

and *r*_*p̄*_ is given by the solution to the equation

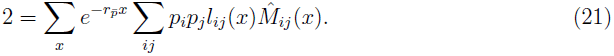

By the same approximation, equation (18) simplifies to

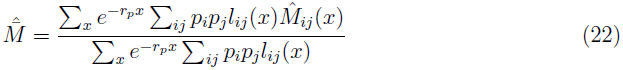

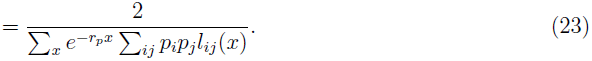

The second line follows from equation (21).

It is common to replace *M*, the number of offspring produced by a female, by *m*, which represents the number of daughters she produces. Assuming equal sex ratios among zygotes, *m* = *M*/2. This can be substituted in to remove a factor of two from each equation. Extending these results to multiple independent loci involves only summing the results across loci. Then the equations simplify to equations (9), (11), (12), and (13) in the text. Equation (19) is, in fact, the covariance between *p*_*i*_ and relative fitness as defined in equation (10).

## Footnotes

1 In dynamical systems theory, the state variables of a model entirely describe the system at any time. We have a dynamically complete model if we can derive the value of state variables at later times, using assumptions about the forces acting on the system.

2 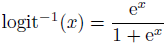

## References

R. Boyd and P. Richerson. Culture and the evolutionary process. University of Chicago Press, 1985.

L. L. Cavalli-Sforza and M. Feldman. Cultural transmission and evolution: a quantitative approach. Number 16. Princeton University Press, 1981.

B. Charlesworth. Evolution in age-structured populations, volume 2. Cambridge University Press Cambridge, 1994.

A. Goodman, I. Koupil, and D. W. Lawson. Low fertility increases descendant socioeconomic position but reduces long-term fitness in a modern post-industrial society. Proceedings of the Royal Society B: Biological Sciences, 279(1746):4342–4351, 2012.

J. H. Jones and R. Bliege Bird. The marginal valuation of fertility. Evolution and Human Behavior, 35(1):65–71, 2014.

P. Käär and J. Jokela. Natural selection on age–specific fertilities in human females: comparison of individual–level fitness measures. Proceedings of the Royal Society of London. Series B: Biological Sciences, 265(1413):2415–2420, 1998.

H. Korpelainen. Human life histories and the demographic transition: a case study from finland, 1870–1949. American journal of physical anthropology, 120(4):384–390, 2003.

J. B. McGraw and H. Caswell. Estimation of individual fitness from life-history data. American Naturalist, pages 47–64, 1996.

J. A. Moorad. A demographic transition altered the strength of selection for fitness and age-specific survival and fertility in a 19th century american population. Evolution, 2012.

J. E. Pettay, L. E. Kruuk, J. Jokela, and V. Lummaa. Heritability and genetic constraints of life-history trait evolution in preindustrial humans. Proceedings of the National Academy of Sciences of the United States of America, 102(8):2838–2843, 2005.

S. C. Stearns, S. G. Byars, D. R. Govindaraju, and D. Ewbank. Measuring selection in contemporary human populations. Nature Reviews Genetics, 11(9):611–622, 2010.

M. Turelli. Phenotypic evolution, constant covariances, and the maintenance of additive variance. Evolution, 42(6):1342–1347, 1988.

M. Turelli and N. Barton. Dynamics of polygenic characters under selection. Theoretical Population Biology, 38(1):1–57, 1990.

M. Turelli and N. Barton. Genetic and statistical analyses of strong selection on polygenic traits: what, me normal? Genetics, 138(3):913–941, 1994.

E. Voland. Differential reproductive success within the krummhærn population (germany, 18th and 19th centuries). Behavioral Ecology and Sociobiology, 26(1):65–72, 1990.

